# Harmonizing single cell 3D genome data with STARK and scNucleome

**DOI:** 10.1101/2025.05.10.653247

**Authors:** Wen-Jie Jiang, KangWen Cai, YuanChen Sun, An Liu, HanWen Zhu, RuiXiang Gao, Chunge Zhong, Nana Wei, Futing Lai, Teng Fei, Yu-Juan Wang, Xiaoqi Zheng, Ming Xu, Hua-Jun Wu

## Abstract

Single-cell three-dimensional genome sequencing (sc3DG-seq) is advancing our understanding of genome regulation and cellular heterogeneity in diverse biological processes. Despite significant technological advancements, a universal tool capable of processing sc3DG-seq data and benchmarking the performance of various techniques in resolving 3D chromatin structures is absent. To fill this gap, we present STARK, a versatile toolkit designed for the preprocessing, quality control and analysis of all spectrum of sc3DG-seq data. Utilizing STARK, we systematically processed 11 sc3DG-seq technologies’ data, enabling a quantitative benchmarking of each technology’s strengths and limitations. Additionally, we developed the EmptyCells algorithm to distinguish high quality from empty barcodes, and introduced the Spatial Structure Capture Efficiency (SSCE) metric to assess the ability of single cells to capture chromatin structures. Furthermore, we established scNucleome, an extensive repository of uniformly processed sc3DG-seq datasets, offering a foundational resource to catalyze further exploration and discovery in the 3D genome research.

## Main

Chromosome conformation capture technology plays a crucial role in elucidating the three-dimensional (3D) genome organization, revealing the complexities of gene expression, regulation, and cellular functions^1-4^. This technique and its derivatives allow us to study the functionality of chromatin domains, chromosomal interactions, and spatial associations between genes and regulatory elements, which are critical for understanding gene regulation during cell differentiation, tissue development and disease progression^5-13^. Despite the contributions of these techniques, their applications to mixed cell populations often result in an aggregate signal that masks cell-to-cell variability and dynamic processes^14, 15^. In light of these limitations, recent technological breakthroughs in single-cell genome-wide mapping of chromatin contacts have significantly advanced our understanding of chromatin organization, particularly in elucidating the dynamics of the cell cycle and revealing the complex organization of tissues.

Currently, over ten methods of sc3DG-seq have been developed, including scHi-C^16^, scHi-C^+17^, sciHi-C^18^, Dip-C^19^, sn-m3C^20^, scMethyl^21^, HiRES ^22^, scSPRTIE^23^, snHi-C^14^, snHi-C^+^, and scNanoHi-C^24^. These methods can be broadly categorized into three types: First, methods that apply bulk Hi-C techniques directly to individual cells post single-cell sorting; second, the use of single-cell high-throughput profiling methods that leverage cellular barcodes; and third, the integration of additional omics, including DNA methylation and gene expression, through simultaneous sequencing within the same single cell. The first type of technology adapts the standard Hi-C protocol^25-27^ by performing ligation after nuclear lysis and dilution of chromatin complexes. This approach processes each cell individually, leading to a more consistent quality and total contacts across single cells. The second type of technology features high-throughput capacity by introducing cellular barcodes at various experimental steps, enabling parallel sequencing of chromatin contacts across a larger cell population. Although variations in total contacts across individual cells may be more pronounced, it significantly expands the scope of analysis for complex tissues. In this category, scSPRITE, an advancement of the SPRITE method^28^, and scNanoHi-C, the first technology to use long-read sequencing in 3D genome analysis^29^, are particularly noteworthy. Both technologies are adept at delineating higher-order chromatin interactions, opening new avenues for studying complex chromosomal structures within single cells. The third type of technology, including HiRES, sn-m3C, and scMethyl, integrates sequencing from additional omics with mapping of 3D chromosome structures. This multi-omics approach demonstrates the interplay between the spatial organization of chromosomes and other molecular context, leading to a deeper insight into complex mechanisms of gene regulation within individual cells^20, 22, 30^.

The diversity of sc3DG-seq has led to a challenge in developing versatile software that can accommodate the varied data formats and analysis requirements from the full spectrum of these methods. Current tools focus on specific downstream analyses, offering functionalities like quality control, imputation, clustering, denoising, and the identification of chromatin loops^31-38^. With the rapid advancement of sc3DG-seq technologies, the throughput has increased dramatically, creating a demand for software capable of handling and analyzing large volumes of sequencing data, akin to the progress seen in single-cell RNA-seq. However, an efficient workflow for processing and filtering high-quality sc3DG-seq data for downstream analysis is still lacking. To fill this gap, we propose STARK—a cutting-edge toolkit for <u>S</u>tructural <u>T</u>opology <u>A</u>nalysis and a <u>R</u>ich <u>K</u>nowledge base. STARK offers a systematic workflow for preprocessing diverse sc3DG-seq data with unified, standardized procedures. It provides robust, versatile quality control measures to ensure the selection of high-quality cells and generates unified single-cell chromatin contact maps alongside the necessary downstream analysis results. STARK represents a significant step forward in addressing the analytical challenges posed by the growing complexity and volume of sc3DG-seq data.

Furthermore, different sc3DG-seq technologies each exhibit unique features and capabilities in mapping the 3D architecture of chromosomes, which can significantly impact the resolution and depth of the data obtained. For instance, variations in sequencing depth and resolution across technologies can lead to different interpretations of genome-wide regulatory interactions and insights into the dynamic changes in chromatin topology under various biological processes^39-41^. Another critical determinant of technological superiority is the ability to reserve chromatin contacts within individual cells^42^. These technological nuances suggest that they may be optimized for specific experimental goals, such as high-resolution mapping, multi-omics integration, or the investigation of complex chromatin interactions. Therefore, it is particularly important to evaluate the characteristics of each technology.

To thoroughly assess the performance and applicability of these technologies, we compiled a comprehensive dataset of sc3DG-seq data and employed STARK for unified preprocessing, quality control and analysis. By minimizing systematic biases, we provided an objective benchmarking of the various methods. We developed a comprehensive set of evaluation indices to quantitatively compare the strengths and limitations of each technology, providing researchers with a clear framework to understand the technological differences, streamline data processing, and inform future experimental design. Additionally, we established scNucleome, a comprehensive publicly accessible repository of uniformly processed sc3DG-seq datasets, encompassing diverse cell lines and tissues across the spectrum of available technologies, serving as a valuable resource for the research community.

## Results

### STARK framework

sc3DG-seq has emerged as an essential tool for understanding the variability in 3D chromatin structure among individual cells. The process encompasses cell sorting, chromosome cross-linking, digestion, ligation, and sequencing, culminating in a map of interactions across the genome. Despite the development of over ten distinct sc3DG-seq techniques, each with unique processing requirements, a unified computational framework has been lacking. This gap poses a significant challenge to the research community, hindering the effective utilization of these extensive and valuable data.

To address this challenge, we propose STARK, a unified framework that conducts preprocessing, quality control and downstream analysis for all current sc3DG-seq data types (Fig.1). STARK encompasses three modules: Preprocess (Fig. 1a), Cell Quality Control (Fig.1b) and Downstream Analysis Fig. 1c).

**Fig. 1.**
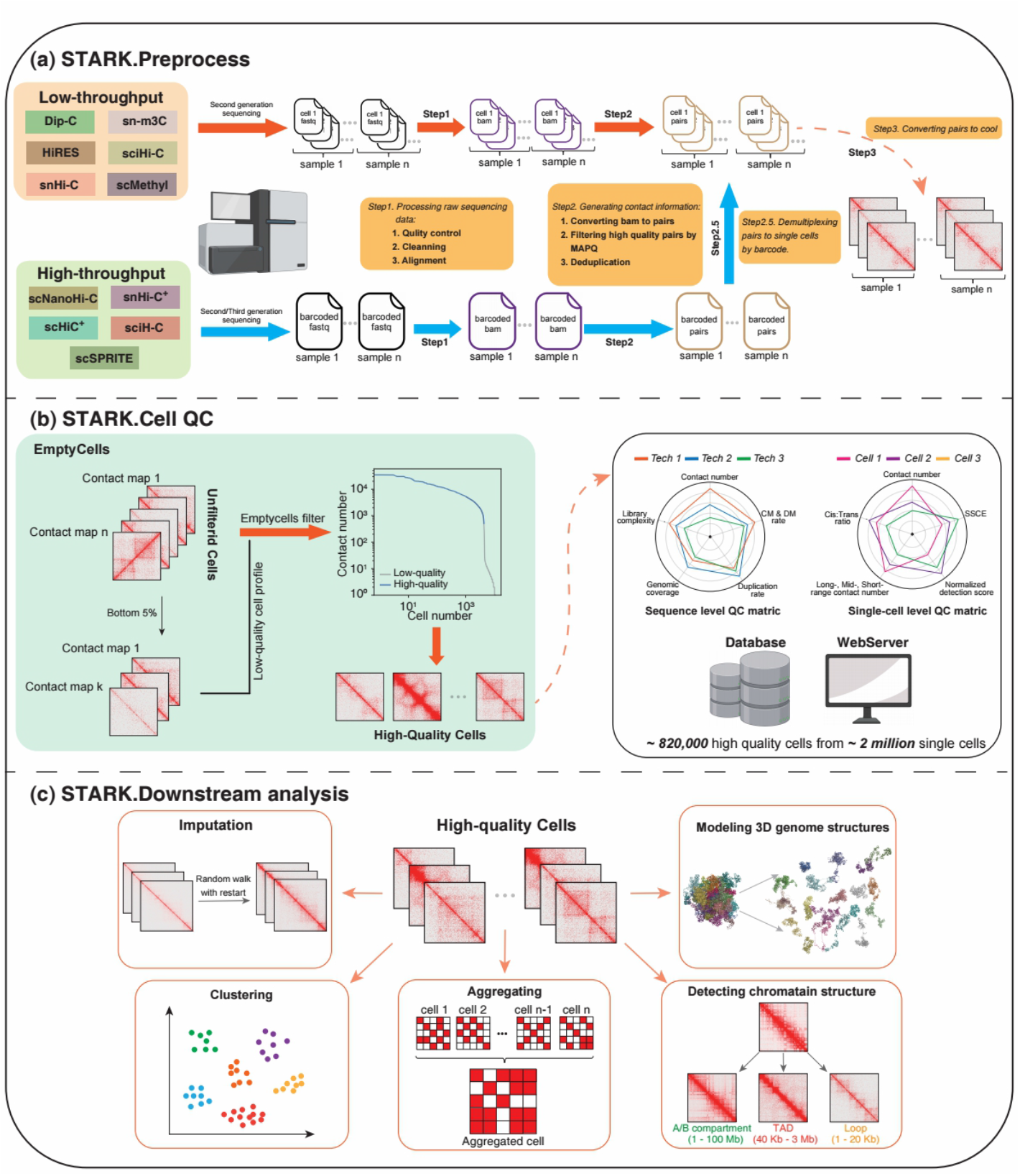
An overview of the STARK workflow. STARK includes preprocess, cell quality control and downstream analysis modules, each designed to work seamlessly with the others to provide a comprehensive pipeline for the analysis of sc3DG-seq data. (a) In STARK.Preprocess module, samples undergo a series of steps including sequencing quality control, cleaning and alignment based on the sequencing platforms used. High-throughput sequencing samples are demultiplexed. The process results in the generation of single-cell contact matrices. (b) STARK.Cell QC module employs the EmptyCells algorithm to filter cells based on contact count, ensuring that only high-quality cells are retained. There cells are then evaluated using a suite of metrics, including contact number, GiniQC, and long-/mid-/short-range contact rates. (c) STARK.Downstream analysis module encompasses a range of analytical processes, including imputing contact matrix, clustering cells, aggregating single cell contact matrices, identifying A/B compartments, TADs, and DNA loops, as well as modeling 3D genome structure.

In Preprocess module (Fig.1a), we consolidate raw data from all techniques into a unified framework. Initially, we extract high-quality sequencing reads and perform sequence alignment using an aligner suited to the sequencing platform. Subsequently, we parse pair information for chromatin interactions, process these based on enzyme cleavage sites, and filter to obtain high-quality chromatin interactions. For techniques employing barcodes, we include a dedicated step for demultiplexing and cell allocation. Finally, we convert processed data into a binary format (.cool) and apply Hi-C correction to each cell.

In Cell Quality Control (QC) module (Fig.1b), we introduce the EmptyCells algorithm, a newly developed approach to filtering and comprehensive quality control of sc3DG-seq data. By first eliminating cells with insufficient interactions and then employing Monte Carlo simulations to identify high-quality cells, EmptyCells ensures that only the most reliable data is used in downstream analyses. To thoroughly evaluate each single cell, we offer a suite of metrics that provide a comprehensive insight into chromatin interactions. These include the total number of contacts, GiniQC, short-range to long-range contact rates, and other relevant measures. Furthermore, we propose the Spatial Structure Capture Efficiency (SSCE), a new metric that assesses a single cell’s ability to capture spatial chromatin structures. The SSCE integrates multiple topological features, enhancing the selection of cells that exhibit informative structural patterns. The SSCE is designed to ensure that cells with fewer overall contacts but notable structural patterns are not disregarded, therefore, complements existing QC metrics.

Downstream Analysis module (Fig.1c) offers a suite of analytical tools that impute, cluster, and aggregate chromatin interactions in single cells, as well as identify key genomic topology such as, A/B compartments, topologically associating domains (TADs) and chromatin loops, and reconstruct 3D genome configurations from 2D interaction maps.

In summary, STARK is a powerful software suite capable of processing extensive datasets from various sc3DG-seq techniques. By processing current and future sc3DG-seq data, STARK empowers the research community to conduct in-depth analyses and unlock new discoveries in the study of 3D genome organization.

**Table 1.**
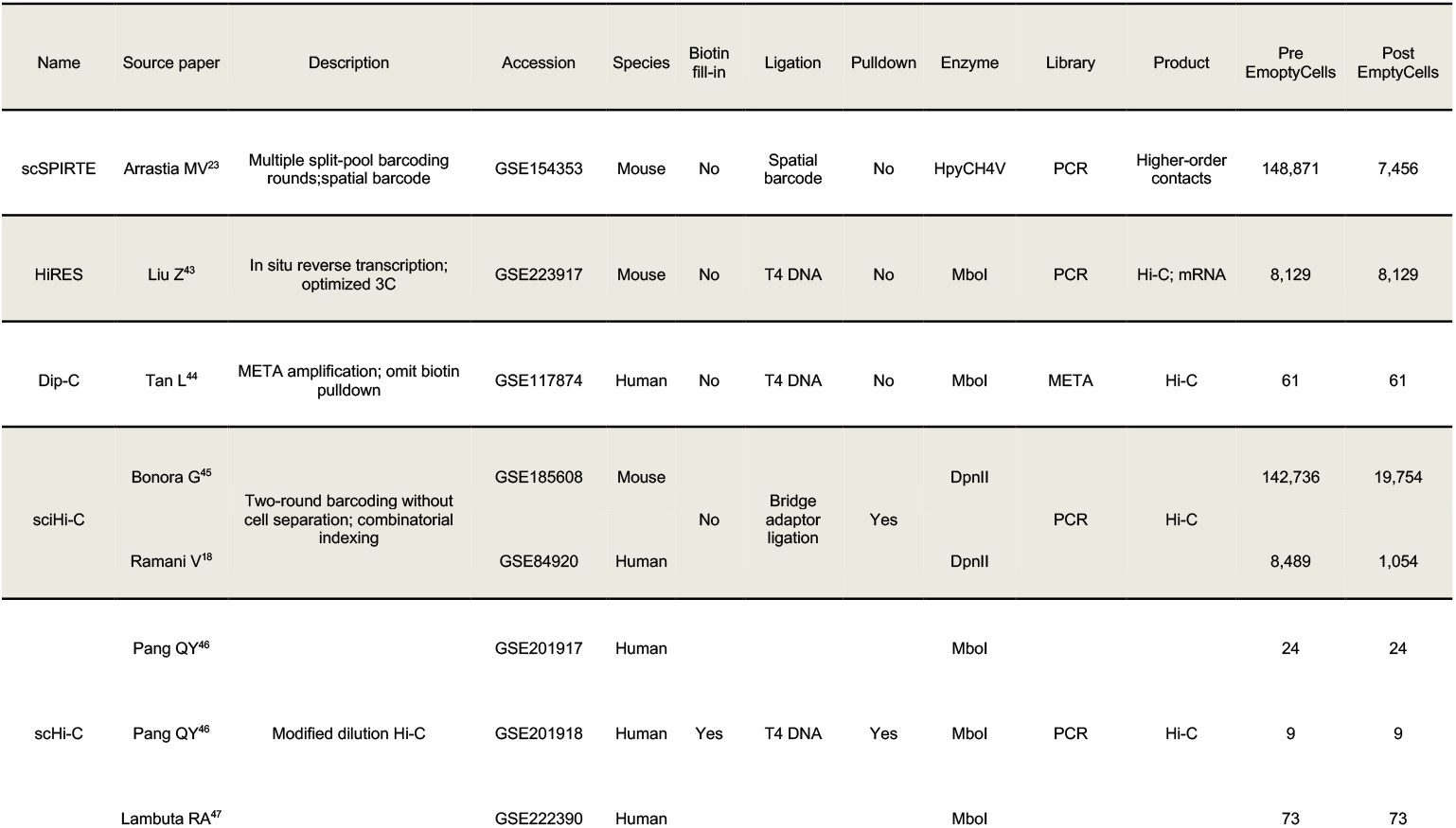
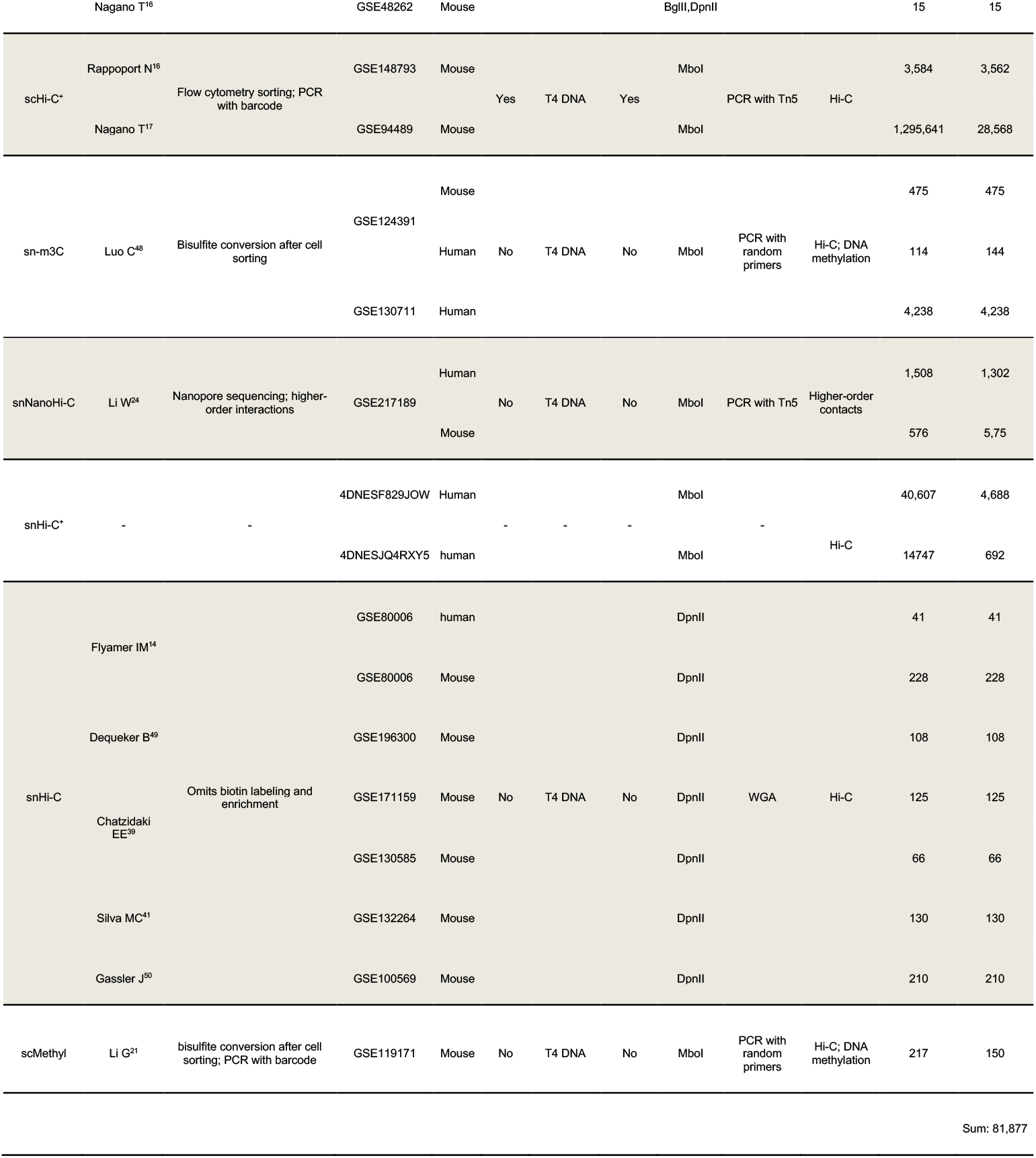
Summary of sc3DG-seq datasets.

### Benchmark on read-level sequencing efficiency

To thoroughly evaluate the current sc3DG-seq technologies, we analyzed each technique data uniformly preprocessed using STARK (see Methods). We began by assessing the sequencing depth and throughput of the various sc3DG-seq technologies (Fig.2a). Low-throughput methods (w/o cell barcode) like snHi-C demonstrate greater sequencing depth, whereas high-throughput methods (with cell barcode), such as snHi-C^+^, sequence a higher number of cells per experiment. In particular, the scSPRITE technology achieves the highest average number of contacts per cell, with its approach to chromatin contact capture being a primary factor. scSPRITE utilizes sonication to fragment the cross-linked DNA complexes, releasing DNA spatial clusters, which are then labeled with spatial barcodes^23, 51^. This method, unlike ligation-based technologies, preserves a broader range of spatially proximate contacts, leading to a unique distribution of captured interactions. The ability of snHi-C to obtain the second highest average number of contacts per cell is largely due to its use of whole-genome amplification, which increases DNA quantity without substantial sequence bias^52^.

**Fig. 2.**
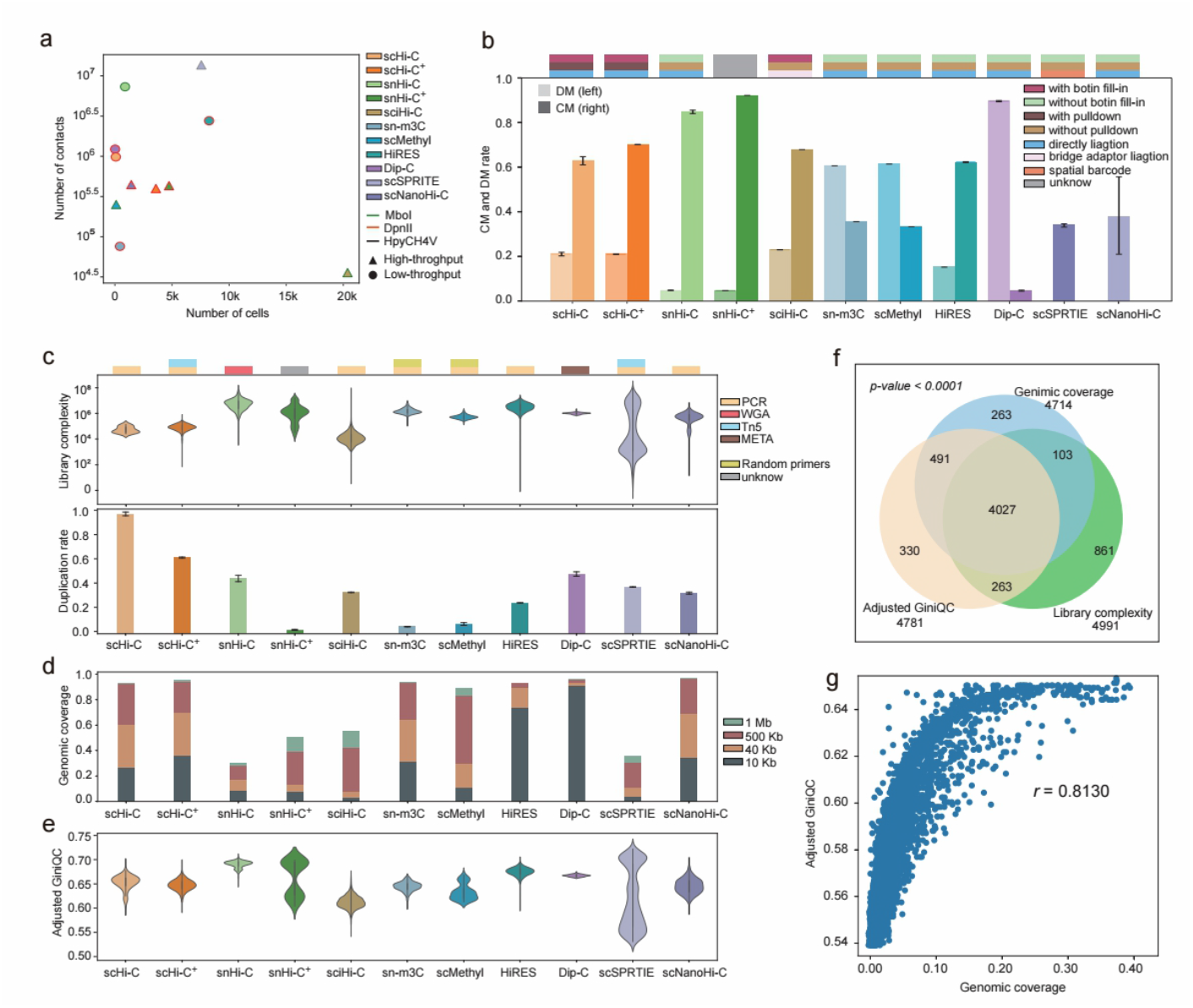
Benchmark of read-level sequencing efficiency across sc3DG-seq technologies. (a) Scatter plot showing the relationship between the number of cells and the average number of contacts per cell. Fill colors of points represent technologies. Border colors of points represent restriction enzymes used. Shapes of points distinguish low- and high-throughput methods. (b) Bar plot displays the rate of concordantly-mapped (CM) and discordantly-mapped (DM) read pairs. Left bars indicate the CM read pairs and right bars indicate DM read pairs. Colors represent different technologies. The bar graph above each pair of bars represents the ligation method, whether biotin is used, and if pulldown is performed. For scSPRITE and scNanoHi-C, only the rate of CM read pairs is calculable. (c) Violin and bar plots represent the library complexity and duplication rate across cells for various technologies, with a corresponding bar above indicating the library preparation method applied. (d) Stacked bar plot demonstrates the average genome coverage across cells at resolutions of 1 Mb, 500 Kb, 40 Kb, and 10 Kb for different technologies. (e) Violin plot exhibits the distribution of GiniQC values across cells for different technologies. (f) Venn diagram illustrates the overlap among cells with low genomic coverage at 1 Mb resolution, low GiniQC and low library complexity in the scSPRITE dataset. (g) Scatter plot depicts the correlation between GiniQC and genome coverage at 1 Mb resolution across the overlapped cells in f.

To evaluate the efficiency of various sc3DG-seq technologies in capturing chromatin contacts and the impact of ligation methods on sequencing outcomes, we calculated the rate of concordantly-mapped (CM) and discordantly-mapped (DM) read pairs (Fig.2b). The majority of technologies demonstrate effective capture capabilities, as evidenced by decent proportion of CM read pairs. However, some methods, such as Dip-C, produce a higher proportion of DM read pairs, possibly due to the use of the META whole-genome amplification approach. The META method is known to reduce chimeric reads^53^, and enhance the quality of CM read pairs (Supplementary Fig.1a), but it may also lead to an increased generation of DM read pairs, suggesting a trade-off in the amplification process. Technologies that simultaneously capture DNA methylation and 3D genome data in the same sets of single cells also show a higher proportion of DM read pairs, potentially due to the bisulfite treatment step required for methylation profiling, which can lead to a decreased alignment rate. In contrast, scSPRITE and scNanoHi-C, which are designed to generate higher-order chromatin contacts, differing from the paired contacts between two genomic regions captured by other methods, are primarily evaluated on their ability to capture these complex interactions effectively, and they also maintain satisfactory alignment rates.

We next evaluated the library complexity and sequence duplication rate across various sc3DG-seq technologies (Fig.2c), revealing significant variability. Low-throughput techniques, which afford more meticulous processing of individual cells, generally result in higher library complexity and lower duplication rate (Supplementary Fig.1b-c). Furthermore, methods using whole-genome amplification (WGA) typically achieve higher library complexity but also came with the trade-off of higher duplication rate (Supplementary Fig.2 d-e).

We then employed the genomic coverage across different resolutions to evaluate the sequencing efficiency of different sc3DG-seq technologies. More than half technologies demonstrate robust performance, achieving approximately full-genome coverage at a resolution of 1 Mb (Fig.2d), whereas snHi-C, snHi-C^+^, sciHi-C, and scSPRITE provide genomic coverage rate ranging from 0.3-0.6. Notably, snHi-C and scSPRITE, despite their lower genomic coverage rate, detected a substantial number of total contacts (Fig.2a). This discrepancy may indicate a propensity for these technologies to preferentially capture compact chromatin structures, a topic we will explore in greater detail in the following analysis.

To assess the sequencing efficiency in an alternative way, we employed the GiniQC metric^54^, which measures the unevenness of read distribution in the contact matrix as an indicator of noise levels. Across the eleven technologies evaluated, average GiniQC values are comparable, suggesting a consistent level of noise. Notably, scSPRITE exhibit a bimodal distribution in genomic coverage (Supplementary Fig.2a), GiniQC (Fig.2e), as well as library complexity (Fig.2c). When analyzing cells with lower genomic coverage, GiniQC, and library complexity, a significant overlap was observed (Fig.2f), indicating a strong correlation between these metrics. The Pearson correlation coefficient between genomic coverage and GiniQC for these cell populations is 0.813 (Fig.2g), highlighting the interdependence of genomic coverage, GiniQC and library complexity in evaluating cell quality. Similar bimodal patterns and results are obtained in scMethyl and snHi-C (Supplementary Fig.2b-2c).

### Benchmark on the efficiency of capturing contacts at various genomic ranges

Chromatin interactions across various genomic distance scales have distinct biological implications (Fig.3a). Short-range interactions, typically less than 20 Kb apart, often denote spatial contacts between local regulatory elements, such as proximal enhancers and promoters, thereby playing a key role in gene transcriptional regulation. Mid-range interactions, ranging from 20 Kb to 2 Mb, indicate spatial contacts between distal regulatory elements and promoters or the organization of functional gene clusters within topologically associating domains (TADs). These interactions represent the functional blocks of gene regulation and insulation, contributing to the precise control of gene transcription^55^. Long-range chromatin interactions, extending over 2 Mb, are essential for the compartment-level folding of chromosomes, influencing transcriptional activity^15, 56, 57^. While less common, inter-chromosomal interactions reflect the spatial organization of chromosomes in the nucleus, sometimes associated with chromosomal translocations and cytogenetic abnormalities^58^. Therefore, a comprehensive understanding of the chromatin interaction landscape across all genomic scales is crucial for elucidating the complexities of gene regulation and delineating the functional organization of the cell nucleus.

**Fig. 3.**
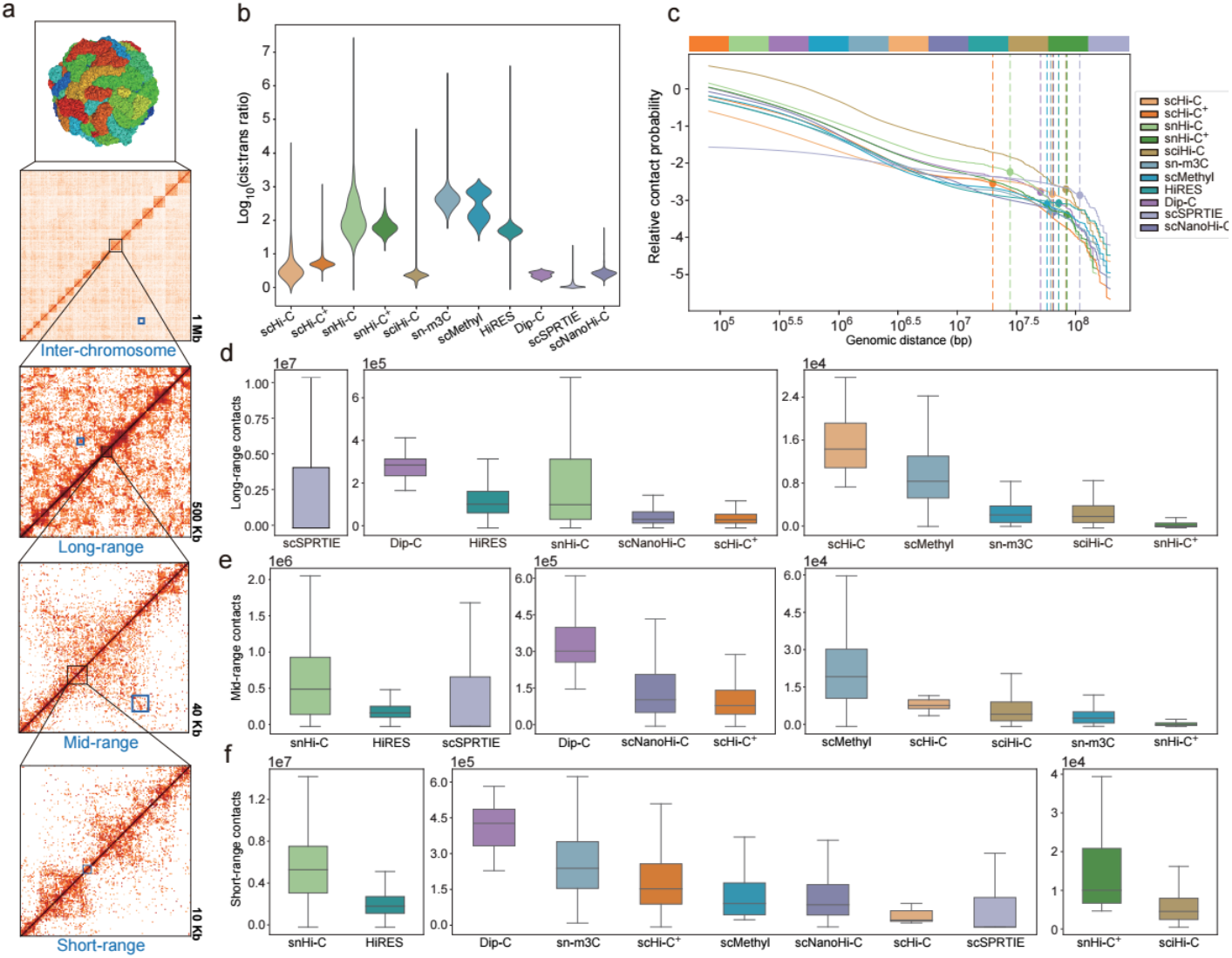
Benchmark of contact capture efficiency across sc3DG-seq technologies. (a) Schematic illustration of contacts at different distance scales and resolutions. From top to bottom: the 3D structure of a cell as obtained by the Dip-C method, contact maps at 1 Mb (with a blue box indicating inter-chromosome contacts), 500 Kb (indicating long-range contacts), 40 Kb (mid-range contacts), and 10 Kb (short-range contacts) resolutions. (b) Violin plot displays the cis:trans ratios for different technologies. (c) Line plot illustrates the relative contact probability against genomic distance for the most deeply sequenced 10 cells across technologies. Points crossing the dashed lines indicates inflection points, with the color bar above the plot showing the order of these points from low to high. (d-f) Box plots depict long-range, mid-range, and short-range contacts, respectively, across technologies. Each is ordered by descending median contact number.

To assess the distance distribution characteristics of chromatin interactions captured by various sc3DG-seq methods, we initially computed the ratio of cis to trans interactions (cis:trans ratio) (Fig.3b). Although all methods detect a higher number of cis interactions, which are interactions between genomic regions on the same chromosome, compared to trans interactions, which occur between different chromosomes, notable variations are observed. snHi-C, snHi-C^+^, sn-m3C, scMethyl, and HiRES yield higher cis:trans ratios than others. Interestingly, the three methods capable of multi-omics sequencing—integrating additional genomic information alongside 3D genome data—consistently demonstrate higher cis:trans ratios. This may be attributed to the more intricate experimental protocols potentially leading to a reduced detection of inter-chromosomal contacts. Additionally, the snHi-C method, which utilizes whole-genome amplification (WGA), also yield favorable cis:trans ratios. It is important to highlight that the average cis:trans ratio for high-throughput methods is higher than those for low-throughput methods (Supplementary Fig.3a).

Among the evaluated methods, scSPRITE exhibit the lowest cis:trans ratio, which is a logical outcome given its use of spatial barcodes. This approach results in a more uniform capture of chromatin contacts across diverse genomic distances. To further investigate this point, we integrated the top 10 cells with the highest number of contacts from each methods and studied the relationship between chromatin interaction frequency and genomic distance (Fig.3c). Consistent with the well-established distance decay law of chromatin interactions in Hi-C data^8^, all methods show a declining trend in interaction frequency with increasing genomic distance. Notably, scSPRITE and scNanoHi-C display a more gradual decline and a prolonged plateau phase, indicating their enhanced ability to capture interactions across a broader range of genomic scales. In contrast, sciHi-C, despite starting with the highest interaction frequency, exhibit the fastest decline, suggesting a greater propensity to capture local chromatin structures at shorter genomic distances. snHi-C, while achieving a favorable cis:trans ratio, show an early inflection point (See methods) in the interaction frequency curve. Coupled with its capacity to detect a large number of contacts, this suggests that snHi-C excels at capturing interactions at shorter genomic distances. The remaining technologies demonstrate relatively uniform performance in capturing chromatin interactions as genomic distance increased.

Finally, we quantitatively compared the number of chromatin interactions captured by different methods across three genomic distance ranges: short-range (< 20 Kb), mid-range (20 Kb - 2 Mb), and long-range (> 2 Mb) (Fig.3d-f). Consistent with our aforementioned findings, scSPRITE outperforms other methods in capturing long-range chromatin contacts. Methods employing whole-genome amplification (WGA)—snHi-C and Dip-C— demonstrate a favorable depth in capturing chromatin contacts across all distance scales, which underscores the substantial benefit of WGA approach. HiRES, which captures both Hi-C and transcriptome, also show a remarkable performance in capturing full range of chromatin contacts. Further analysis of the correlations between the three distance ranges revealed that most methods exhibit higher correlations in the number of captured contacts between adjacent distance ranges (Supplementary Fig.3b).

### Benchmark on the efficiency of capturing topological structures

Chromatin structures play a crucial role in regulating gene expression and orchestrate the functional organization of the cell nucleus. sc3DG-seq serves as a powerful tool for investigating the heterogeneity of chromatin structures among individual cells, with the resolution of detected structures varying^15^. For instance, at a resolution of 1 Mb, chromatin territories can be identified, A/B compartments are detectable at 500 Kb resolution, and topologically associating domains (TADs) become discernable at 40-50 Kb (Fig.4a). Due to the sparsity of sc3DG-seq data, loop detection often requires data enhancement, therefore, we limited our comparison to the original contact maps.

**Fig. 4.**
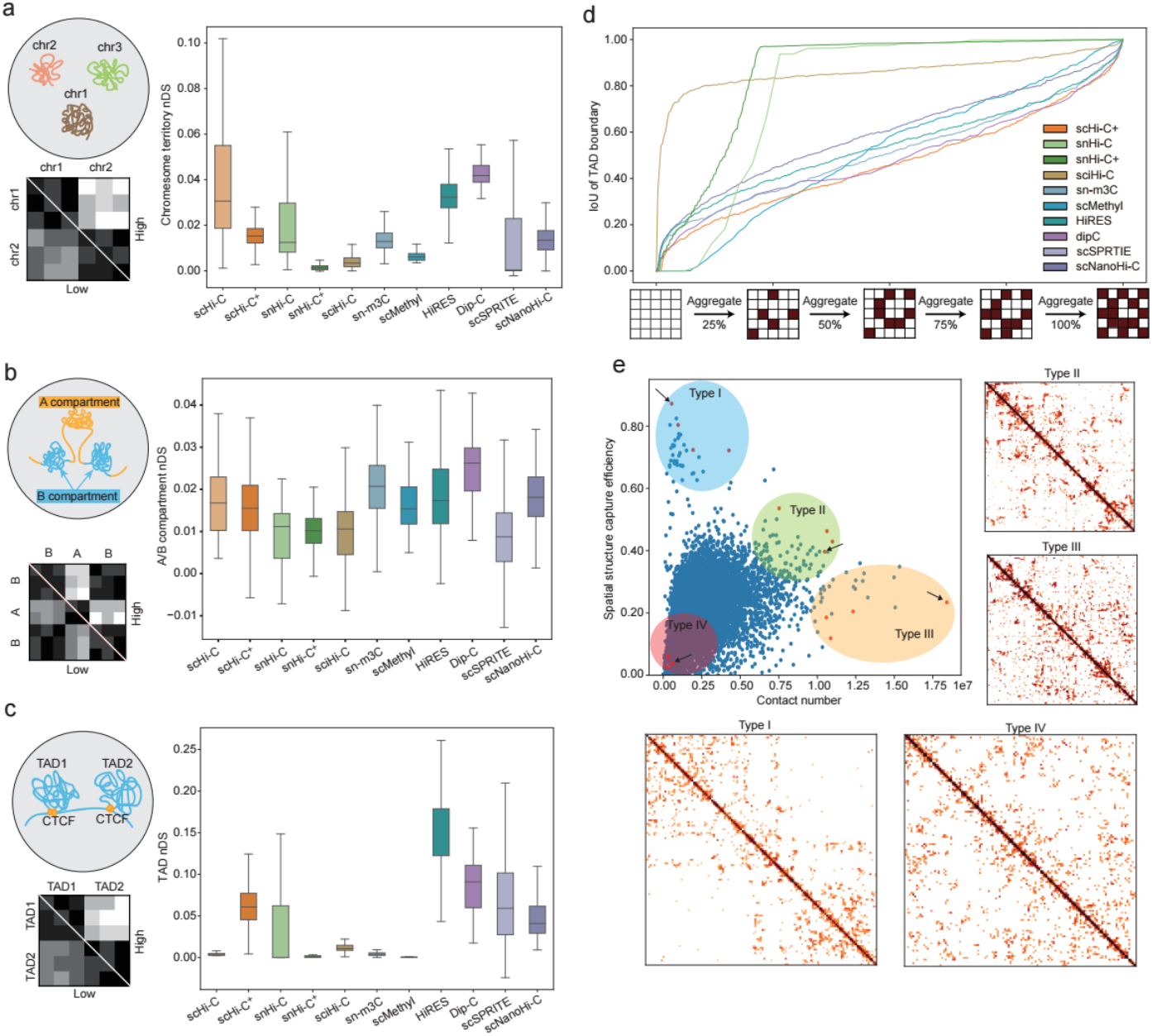
Benchmark of structure detection ability across sc3DG-seq technologies. (a-c) Normalized Detection Score (nDS) of chromosome territory, A/B compartment, and TAD for the top 300 deeply sequenced cells across various technologies. An inset on the top left provides a schematic illustration of these genomic structures, and the diagram on the bottom left shows the pattern of contact maps with high and low nDSs. (d) Line plot displays the cumulative similarity of TAD boundaries between those obtained from aggregated single cells and total cells for different technologies. This analysis is performed on the top 300 deeply sequenced cells in each technology. The y-axis represents the Intersection over Union (IoU) as a measure of similarity. The x-axis indicates the percentage of cells integrated, with a schematic illustration of the integration process shown. (e) Scatter plot demonstrates the relationship between the SSCE and contact number using HiRES data as an example. Cells are categorized as type I-IV, color-coded in blue, green, orange, and red respectively. Each dot represents an individual cell, with red dots providing visualized examples. Heatmaps correspond to the cells indicated by the black arrows in the scatter plot, showcasing the contact map for the region Chr10: 2 Mb-133 Mb. Additional examples are provided in supplementary figures.

To evaluate the ability of different methods to detect chromatin structures, we calculated normalized Detection Scores (nDS)^59^ for single cells derived from each method. To minimize potential biases and enhance computational efficiency, we have revised the original calculation approach (see Methods). Using this improved methodology, we assessed the capability of the 11 sc3DG-seq methods to reconstruct 3D chromatin structures at various resolutions, focusing on chromatin territories, A/B compartments, and TADs. The results indicate that most technologies can detect these structures to some extent (Fig.4a-c). In the detection of chromatin territories, low-throughput methods generally yield higher nDS compared to high-throughput methods (Supplementary Fig.3c). This may be due to the washing steps involved in barcoding of single cells, which can lead to the loss of detection efficiency of chromatin territories. This aligns with our cis:trans ratio analysis findings. Similarly, for A/B compartment detection, low-throughput methods outperform high-throughput methods (Supplementary Fig.3d). However, in discerning TAD structures, high-throughput methods slightly surpass low-throughput methods (Supplementary Fig.3e). We hypothesize that the cell barcoding process may enhance the capture of more compact chromatin structures, which could explain this contrasting observation in TAD detection.

When examining the performance of individual technologies, scSPRITE, Dip-C, scNanoHi-C, scHi-C^+^, and snHi-C consistently achieve higher nDS across various genomic resolutions, with particularly notable advantages in detecting TADs. Their superior performance is likely attributed to their proficiency in capturing mid- and long-range chromatin interactions, as previously discussed in our analysis. Interestingly, multi-omics methods that simultaneously sequencing DNA methylation, which are sn-m3C and scMethyl, demonstrated elevated nDS at the A/B compartment and chromatin territory levels, but lower nDS at the TAD levels. This observation suggests that the incorporation of methylation information may affect the detection of compact chromatin structures.

Next, we examined the ability to reconstruct bulk-like TAD boundaries by integrating an increasing number of single-cell data for different sc3DG-seq technologies (Fig.4d). To achieve bulk-like TAD boundaries, we consolidated the top 300 most abundant single-cell data from each technology. Utilizing the Accumulative Single-cell Quality Analysis method (see Methods), we tracked the trend in similarity between these bulk-like TAD boundaries and those reconstructed from integrating an increasing number of cells. We employed the intersection over union (IoU) as a metric for consistency evaluation. Overall, the consistency scores of all technologies rise with an increasing number of cells integrated, yet there are notable differences in the rate of score increase. snHi-C, snHi-C^+^, and sciHi-C exhibited rapid increase in consistency score. Particularly, sciHi-C reached saturation early on, suggesting its ability to approximate bulk-like characteristics with a small subset of cells. In contrast, other technologies showed a slower progression toward the fitting trend, indicating that they require a larger number of cells to reliably reconstruct a bulk-like TAD structures. These findings imply that TAD structures may be inadequately captured and can be considerably variable among individual cells, highlighting the need for further optimization of experimental protocols to enhance data stability. Notably, some of these variations may also stem from inherent differences between distinct tissues and cell lines, which could influence the observed data consistency.

nDS can capture structure-specific information but require analyses at various resolutions for a comprehensive assessment. To address this, we developed the Spatial Structure Capture Efficiency (SSCE), a new metric that quantitatively evaluates a single cell’s ability to capture spatial chromatin structures (Fig.5e). The SSCE incorporates multiple topological information, including chromatin territories, A/B compartments and TADs, and evaluates their contribution to the total contacts within each cell. Based on the SSCE and total contacts, we categorized cells into four distinct types I - IV. Notably, Type I cells, which have a relatively high SSCE but low total contacts, are observed to exhibit more pronounced structures despite similar sequencing depth as Type IV cells (Fig.5e and Supplementary Fig.4a-b). Conversely, Type II and Type III cells, while having a relatively high sequencing depth, show lower SSCE values, and their contact maps indicate a higher, albeit structural capturing capability, noise. This holistic approach ensures that cells with fewer overall contacts but significant structural patterns are not overlooked. Traditional quality control methods that rely solely on total contacts may inadvertently exclude cells with lower sequencing depths that still capture important structures. By incorporating SSCE, we can enhance the retention of cells that capture critical structures. Therefore, SSCE serves as a valuable complement to existing quality control measures for sc3DG-seq data, aiding in the downstream selection of single cells for further analysis.

**Fig. 5.**
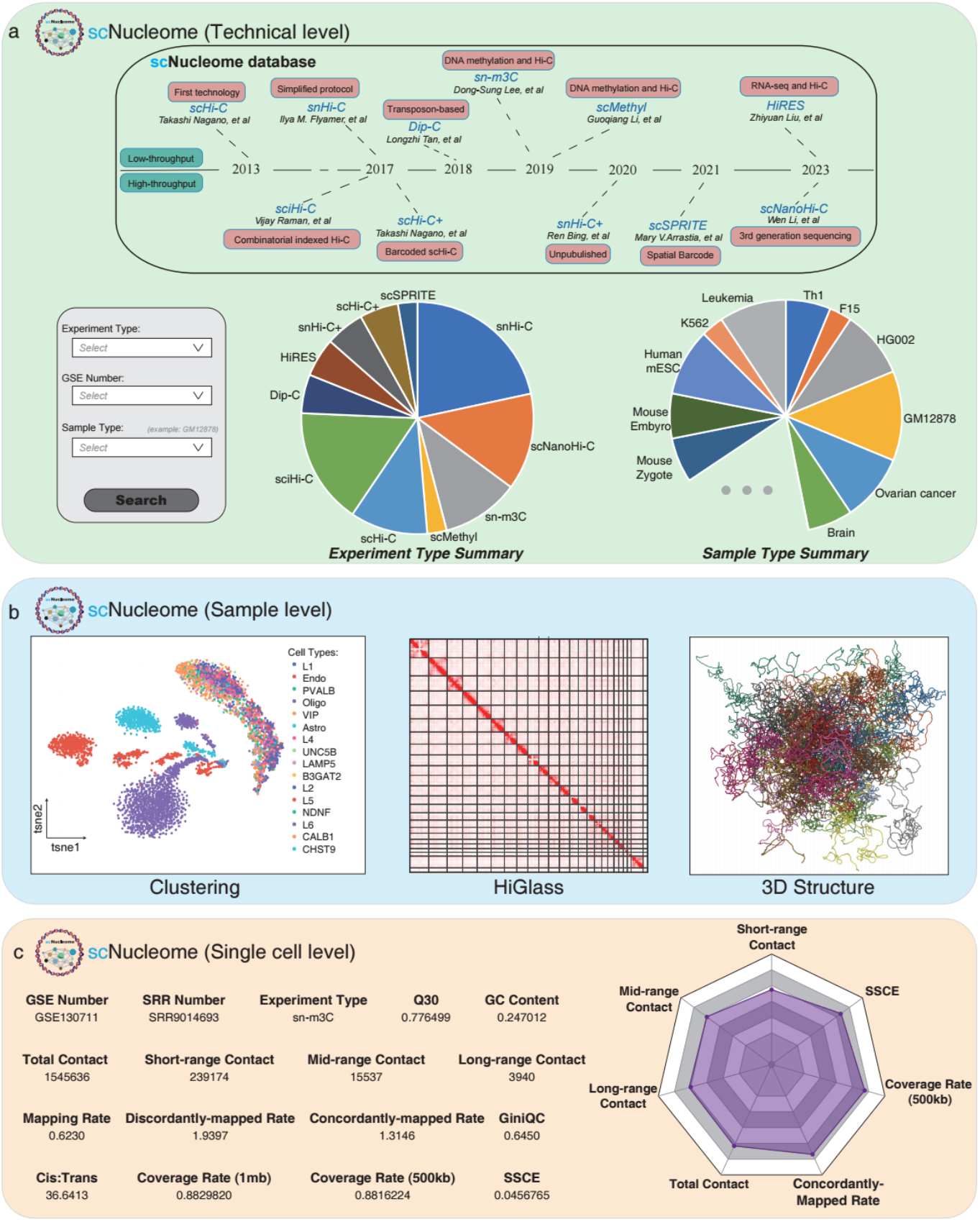
Overview of the scNucleome database. (a) Milestones in the development of sc3DG-seq technologies (upper panel) and summary of the sc3DG-seq data collected in the database (lower panel). (b) Example of the analyses results and visualization choices available within the database. (c) Example of a quality control report (left) and radar visualization (right) for an individual cell available in the database, which offers a multi-dimensional view of the cell’s characteristics and quality metrics.

### scNucleome: a harmonized sc3DG-seq database

Processing large-scale sc3DG-seq data from scratch under a unified framework remains a major challenge due to the high costs of computational resources. To fill this gap, we employed STARK to process approximately 2 million single cells from 24 sc3DG-seq datasets, encompassing 40 sample types. This extensive effort has resulted in scNucleome, currently the most comprehensive single-cell 3D genome database (Fig.5a).

scNucleome facilitates data search by technology, sample type, and data source, enabling precise and efficient data retrieval. For each dataset, we provide multi-dimensional data analysis results, including low-dimensional representation, cell clustering and annotation, contact map visualization by HiGlass^60^, and simulated 3D organization model by Mol* Viewer^61^, allowing researchers to explore datasets from various perspectives (Fig.5b). Furthermore, scNucleome offers a wide array of quality control metrics at both the dataset and single-cell levels, including library complexity, CM/DM read pair rates, total contacts, GiniQC, nDS, SSCE, and more. These metrics are designed to aid in further experimental design, benchmarking and customizable cell filtration (Fig.5c). We anticipate that this comprehensive platform will markedly accelerate research progress, providing a robust foundation for advancements in the field of 3D genome studies.

## Discussion

In this study, we introduce STARK, a comprehensive computational framework designed to provide a unified and standardized approach for processing and analyzing a wide array of sc3DG-seq data. STARK encompasses three core modules - Preprocess, Cell QC, and Downstream Analysis – enabling unified data handling, extensive quality control, and an in-depth exploration of chromatin structures within individual cells.

A key strength of STARK lies in its adaptability to the diverse data formats and analytical requirements across the full spectrum of current sc3DG-seq technologies. This includes capabilities for processing data from both low and high-throughput technologies, as well as accommodating second and third-generation sequencing-based methods and single and multi-omics approaches. In addition, STARK introduces EmptyCells, a new quality control method tailored for high-throughput technologies, and proposes the SSCE, a new metric for evaluating the capture efficiency of chromatin structure in single cells. By systematically processing approximately ∼2 million single cells across eleven different sc3DG-seq techniques, STARK offers a comprehensive benchmarking of the strengths and limitations inherent to each method. This comprehensive assessment empowers researchers to make informed choices regarding the most suitable technology for their experimental objectives, whether they aim for high-through in cell number, high-resolution contact mapping, or the investigation of complex chromatin interactions.

A limitation of the current STARK framework is its focus on processing single-cell chromosome conformation data, without the support for integrating multi-omics information. Emerging sc3DG-seq technologies, including HiRES, sn-m3C, and scMethyl, enable the simultaneous capture of chromatin contacts and additional genomic information, such as DNA methylation or gene expression, within the same single cells. Extending STARK’s functionality to incorporate such multi-omics information could further enhance its utility, enabling researchers to gain deeper insights into the interplay between chromatin architecture and other molecular processes.

Additionally, while STARK has demonstrated comprehensive capabilities in processing and quality control for sc3DG-seq data, the current analytical functions it offers are relatively basic. Incorporating STARK with a broader range of advanced analytical features, such as data imputation, integration and clustering, could enhance its value as a one-stop solution for comprehensive sc3DG-seq data exploration.

Despite these limitations, the STARK framework represents a significant step forward in addressing the analytical challenges posed by the growing complexity and volume of sc3DG-seq data. By providing a unified, standardized, and robust platform for data processing and quality control, STARK empowered the research community to conduct in-depth analyses and unlock new discoveries in the field of 3D genome organization. Furthermore, the introduction of scNucleome, a publicly accessible repository featuring a curated collection of uniformly processed sc3DG-seq datasets, is poised to catalyze collaborative efforts and accelerate progress in this rapidly evolving field.

## METHOD

### STARK Framework

The STARK Framework is a comprehensive software suite designed to manage the entire analysis workflow for single-cell Hi-C data. It seamlessly integrates three main functions, each tailored to address specific stages of the workflow:

STARK is a software suite designed to manage the complete analysis workflow for sc3DG-seq data. It comprises three main modules, each tailored to address specific stages of the workflow: (a) STARK.Preprocess, (b) STARK.Cell QC, and (c) STARK.Downstream Analysis. The following sections are the description of the implementation:

(a) STARK.Preprocess:

We initiate the preprocessing by performing sequence quality control and adaptor trimming on the raw fastq files from sc3DG-seq data using fastp^62^ with default parameters. This step yields trimmed fastq files. For technologies that do not involve bisulfite conversion, the trimmed fastq files are aligned to the reference genome (hg38/mm10) using one of the aligners: BWA, Bowtie2, or minimap2^63-65^, selected based on the characteristics of the sequencing technology. For sn-m3C and scMethyl technologies, a two-round alignment approach is employed. The first-round alignment is conducted using bismark^66^ to map the fastq files to the reference genome. Sequences not aligning in the first round are trimmed by 40-bp from both ends, and a second alignment round is performed using the same settings. The alignment results are stored into BAM format and are subsequently filtered and sorted using samtools^67^.

Next, we proceed to parse the BAM files to generate “pairs” files, we apply stringent filtering criteria to ensure high-quality pairs: only pairs with a mapping quality score (mapq) of 30 or higher for both fragments are retained. Subsequently, we use the STARK to generate whole-genome enzyme cutting sites for respective experimental enzymes and assign the nearest restriction sites to each alignment in a pair. For high-throughput technologies, including scNanoHi-C, snHi-C^+^, scHi-C, scHi-C^+^, and scSPRITE, we extract cell barcode from the raw sequencing files, linking each read to its originating single cell. Using cell barcodes, we demultiplex the contacts in the “pairs” files down to the single-cell level. The “pairs” files are then sorted, deduplicated and filtered to eliminate PCR duplicates and other artifacts. Finally, we convert the “pairs” format to the “cool” format, correct the contact matrix, and generate single-cell multi-resolution “cool” files.

(b) STARK.Cell QC

Quality control on cell level is a critical step to filter out low-quality single cells in high-throughput sequencing data analysis. We develop the “EmptyCells” algorithm to perform this task (details provided below). Initially, the algorithm removes single cells with fewer than T contacts. It then models the bottom 5% of cells with lowest contacts and applies a Monte Carlo simulation method to test each cell, identifying and removing those that are significantly low-quality. Cells that pass this filtering are considered to have high-quality single-cell 3D structures.

Next, we implement a wide range of quality control (QC) metrics to provide a comprehensive assessment for each single cell. These QC metrics include the total contact number, genomic coverage, the rates of short-range (< 20 Kb), mid-range (20 kb - 2 Mb), and long-range (> 2 Mb) contacts, GiniQC, normalized Detection Scores (nDS), and a newly developed metric – the Spatial Structure Capture Efficiency (SSCE). The SSCE is designed for a more nuanced assessment of the structural capture efficiency in single cells. These metrics are discussed in greater detail in the corresponding section of the Methods.

(c) STARK.Downstream Analysis

#### 1. Imputation via Random Walk with Restart (RWR)

The RWR algorithm is a variant of the traditional random walk technique. It is designed to explore the graph by randomly walking through its nodes, with the added feature that at each step, there is a probability the walk will “restart” and return to a predefined starting node. We implement this algorithm, as referenced in previous research^68^, for the interpolation of single-cell 3D genome data.

First, we construct a probability transition matrix (*P*) by normalizing the single-cell contact matrix (*A*) such that each row sums to 1. The walk restarts at the starting node with a probability *r*, and with the probability 1−*r*, it proceeds to a neighboring node. The goal is to find the steady-state *π* which represents the matrix given the restart probability *r*. The RWR is solved iteratively through the following steps:

1. Initialization: Start with an initial probability vector *π*^0^ = *A*.
2. Iteration: Update *π* using the formula: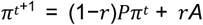. *P* is the transition probability matrix.
3. Convergence: Repeat the iteration until π converges, when 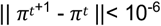.

Upon completion of the iterations, we obtain 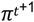 which serves as the imputated single-cell contact matrix.

#### 2. Dimensionality reduction and clustering

sc3DG-seq data is typically represented as a two-dimensional (2D) contact matrix for each chromosome. However, this format is not conducive to dimensionality reduction processes. To address this, we flatten each 2D matrix into a one-dimensional (1D) vector, and then concatenate these vectors to create a long vector representative of each cell’s data. We proceed by normalizing the data with respect to library size to adjust for differences in sequencing depth. For dimensionality reduction, we apply Principal Component Analysis (PCA) to the normalized data. This allows us to extract the primary principal components (PCs) that capture the most variance within the dataset. Subsequently, we utilize Uniform Manifold Approximation and Projection (UMAP)^69^ to transform the PCs into a two-dimensional (2D) representation. Visualization on this 2D space facilitates the identification of patterns and structures within the data. For the clustering analysis, we employ the Leiden community detection algorithm, a method widely recognized for its effectiveness in the analysis of single-cell RNA sequencing (scRNA-seq) data. This algorithm helps to identify clusters within the single-cell data, providing insights into the cellular heterogeneity and potential cell types or states.

#### 3. Cell aggregation

Given the sparsity of sc3DG-seq data, an integrated analysis of data from multiple cells is often beneficial. STARK facilitates this by enabling the aggregation of data from multiple single cells for subsequent analysis. The aggregation is mathematically represented by the equation:

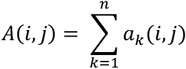

Matrix A denotes the integrated 3D genome data from multiple cells, and *a*_*k*_ represents the single-cell 3D genome data from the k-th cell. The indices i and j indicate the genomic positions of interactions. This aggregation function serves to generate a pseudo-bulk dataset that represents cell types or clusters of interest. The creation of pseudo-bulk data allows for the enhancement of signal-to-noise ratio and can reveal patterns that may not be evident when analyzing single cells in isolation. This approach is particularly useful for downstream analyses such as identifying consensus chromosomal structures, studying cell type–specific interactions, and exploring the organization of the genome across different cellular states.

#### 4. Detecting chromatin structure

Analyzing 3D genome sequencing data to identify A/B compartments, Topologically Associating Domains (TADs), and chromatin loops is crucial for understanding the spatial organization of chromatin and its role in gene expression and regulation. Below are the methods we implemented for identifying these three structural features in STARK:

A/B Compartments: We begin by conducting Principal Component Analysis (PCA) on preprocessed contact matrices. PCA helps capture interaction patterns between chromatin regions and projects them into the principal component space. The first principal component (PC1) is analyzed for positive and negative values by gene density, allowing us to distinguish A compartments, which are associated with active regions, from B compartments, which are inactive.

TAD Identification: We apply the Insulation Score algorithm^70^ to the contact matrix. This score detects TAD boundaries by calculating the average change in contact frequency within sliding windows along the chromatin sequence. Significant increases in Insulation Score values indicate TAD boundaries.

Loop Identification: The process of identifying loops involves pinpointing significant hotspots in the contact matrix, which represent frequent interactions between distal genomic locations. We employ peak calling methods^70^ to identify these loops, using statistical analysis to filter hotspots in the contact matrix and identify regions with significantly higher interaction frequencies compared to the background.

These methods provide a comprehensive approach to detecting key chromatin structures that are vital for elucidating the complex interactions within the genome.

#### 5. Modeling 3D-genome structures

Reconstructing the 3D structure of chromatin from a contact matrix involves several steps. Initially, the contact matrix undergoes preprocessing to eliminate noise and adjust its resolution. Following this, distance constraints are derived from the matrix, which serve as the foundation for establishing the 3D structure. Then, Monte Carlo simulation is employed to find chromatin structures that align with these constraints. We refer to the nuc_dynamics^71^ for the code implementation of this function.

## EmptyCells

sc3DG-seq technologies are broadly categorized into two approaches: low-throughput non-barcoded and high-throughput barcoded methods. Non-barcoded methods involve the precise isolation of individual cells, which are then subjected to library construction and sequencing. In contrast, barcoded methods allow for the high-throughput sequencing of multiple cells simultaneously by incorporating cell barcodes. While barcoded methods enable the sequencing of a larger number of cells per experiment, it can result in a substantial variation in the number of contacts captured from each cell. Therefore, the effective filtering of low-quality cells is critical. Common quality control approach rely on total contact number thresholds to filter cells/barcodes. However, this approach have limitation, as library preparation efficiencies can vary among barcodes. Moreover, stringent thresholds may inadvertently exclude cells with less compact chromatin structures, which could be particularly problematic when analyzing cell-state transitions^17^.

Inspired by EmptyDrops^72^ and Cellranger (10 x Genomics, version 7.0.1), We propose a two-step method for filtering low-quality cells in sc3DG-seq data: (i) Preliminary filtering by setting a low threshold based on the total number of contacts to remove low-quality cells with insufficient sequencing or capture efficiency. (ii) Further refinement by modeling the profile of these low-quality cells to enhance the filtering process. We hypothesize that high-quality cells exhibit a diverse array of interaction types. Therefore, we incorporate the variety of interactions between different chromosomes as a key feature in our model. Mathematically, all barcodes of a sample can be represented as A = y_bg_∈ R^N×ℊ^ , where *y*_*bg*_ denotes the number of contacts for interaction type g in barcode b, N denotes the total number of barcodes, and ℊ denotes the interaction type. Notably, we exclude interaction types that are not present in all barcodes from our analysis to maintain consistency and accuracy in the filtering process.

In the first step, we establish a threshold T based on the total number of contacts per barcode. Barcodes that have fewer contacts than T are considered incomplete or of low quality and are therefore removed. Importantly, a barcode exceeding the threshold T does not necessarily indicate a complete cell representation; it may still only capture a subset of contacts from a cell. We define the threshold T as follows:

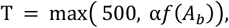

where α is a hyperparameter, which we set to 0.01, f(·) calculates the median value of the barcode contact vector *A*_*b*_, which is represented as the sum of contacts across all interaction types per barcode:

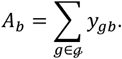

Barcodes that fall below this threshold T are discarded, while the remaining barcodes proceed to the next step of the filtering process.

To construct the profile of low-quality barcodes, we selected a subset of barcodes characterized by a low number of contacts, specifically selecting the bottom 5% to form a set we designate as G. We then calculated A as the sum of contacts across all barcodes in G:

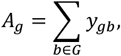

yielding a contact vector A = (*A*_1_, … , *A*_ℊ_).

We apply the Simple Good-Turing algorithm to *A*_*g*_ to obtain the posterior expectation 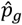 , representing the probability of sampling a contact from an interaction type g. This method ensures that even contact types with zero observed contacts are assigned non-zero probabilities, thereby preventing undefined values in subsequent calculations^73^.

For a given barcode b, the total number of contacts is denoted as *t*_*b*_ and can be modeled using a Dirichlet multinomial distribution. The likelihood of obtaining contacts from barcode b conditional on *t*_*b*_, is defined as follows:

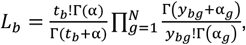

where 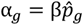 is a scaling factor used to assess the importance of 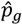 in sampling from A. By modeling overdispersion in contact types through the estimation of β, our model accounts for broader range of variability in contact numbers than a simple multinomial model would allow. For example, a lower β value signifies a greater dispersion level in contact type, which could be due to variations in capture efficiency or differences in library amplification. This approach allows for more accurate p-value calculation when evaluating if a barcode deviates significantly from the low-quality set.

We employ the Monte Carlo simulation method to calculate the p-value for each barcode. For each iteration i of a given barcode, a new contact number vector is randomly sampled. The likelihood *L* ^′^_*bi*_ is then calculated based on the posterior expectation 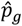,the corresponding total number of contacts *t*_*b*_ and the scaling factor β. The count *R*_*b*_ of likelihoods from the R round simulation that are less than *L*_*b*_ is utilized to compute the p-value^74^:

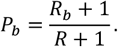

This approach allows us to quantitatively assess the quality of each barcode and to filter out those that do not meet the predetermined quality thresholds, thereby enhancing the reliability of downstream analyses.

## SSCE

Spatial Structure Capture Efficiency (SSCE) is introduced as a quantitative metric designed to evaluate the efficacy of single-cell sequencing in capturing the chromatin structural information. It is crucial to recognize that while the total number of sequenced contacts can vary significantly among cells, a lower count does not necessarily indicate inferior sequencing quality. Such a result may simply reflect the capture of a specific subset of the chromatin architecture, which can still have substantial biological implications. Conversely, a high number of detected contacts does not guarantee the biological relevance of these interactions or confirm an accurate representation of the cell’s chromatin spatial organization. With this in mind, we introduce a novel indicator, termed Spatial Structure Capture Efficiency (SSCE), which quantifies the proportion of intra-cellular contacts contributing to the formation of the cell’s chromatin architecture. This metric is essential for a more nuanced understanding of the structural fidelity of single-cell sequencing data. The SSCE are calculated as follows:

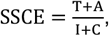

where T, A, I, and B are derived from fitting the following regression:

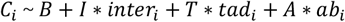

In this equation, *C*_*i*_ denotes the aggregate count of contacts associated with chromosome i, inter_*i*_ represents the number of interactions that chromosome i engages in with other chromosomes in the genome. ta*d*_*i*_ corresponds to the count of TAD boundaries identified on chromosome i at a resolution of 40 kilobases, which is pivotal for elucidating chromatin domain organization. a*b*_*i*_ signifies the count of A/B compartment switches for chromosome i determined at a resolution of 100 kilobases, providing insights into the chromatin’s compartmentalization. The constant B in the equation serves two critical roles. Firstly, it accounts for contacts that cannot be attributed to any specific chromosomal structure feature. Secondly, it includes contacts that may result from technical or systematic errors during the sequencing process. To ensure the accuracy of the model, any chromosomes where all variables are nullified are systematically excluded from the regression analysis.

Considering the inherent collinearity between the counts of a*b*_*i*_ and tad_*i*_, we employ Ridge regression as our fitting method. The coefficients derived from the Ridge regression model are then utilized to compute the SSCE. It is logical to infer that higher values of structural features a*b*_*i*_ and tad_*i*_, along with lower values of non-structural features B and I, contribute to higher SSCE scores. An elevated SSCE score, therefore, implies a more thorough and precise depiction of the single-cell chromatin structure. Additionally, we have confirmed that the loss of contacts due to various factors can result in a decrease in SSCE (Supplemetary Fig.4b).

### GiniQC and genomic coverage

We employ GiniQC^54^ to quantify the internal noise in sc3DG-seq data. GiniQC is also utilized within the STARK framework for data quality assessment. Genomic coverage is determined based on a deduplicated contact matrix, providing an essential measure for evaluating the distribution and comprehensiveness of the data. To thoroughly dissect the genomic interaction landscape, we compute genomic coverage at three resolutions: 10 Kb, 40 Kb, and 100 Kb. These varying resolutions allow for a multi-scale analysis, enhancing our understanding of the 3D genome organization of the genome.

### Duplication rate and library complexity

The duplication rate and library complexity were assessed using ‘pairtools dedup’. To effectively remove PCR duplicates, we utilized the ‘--max-dist 3’ parameter during the deduplication process. The library complexity estimate was derived from the intrinsic estimation method implemented within pairtools. This metric provides a proxy for the number of unique molecular identifiers (UMIs) expected in a hypothetical, duplicate-free library. For the purpose of visualizing and comparing these estimates across various datasets, we applied a log-normalization transformation.

### Normalized detection scores for 3D genome structures

Normalized Detection Scores (nDS) are utilized to evaluate the clarity of 3D genome structures, including chromosome territories, A/B compartments, and TADs. We have refined the approach introduced in scSPRITE, leveraging the high-throughput capabilities of single-cell sequencing while bypassing the requirement for pre-computed genome-wide 3D structural data. The Detection Scores (DS) are computed using the native contact matrix, which defines the frequency of interactions among various genomic bins. A higher DS for chromatin compartments in a cell, for instance, suggests a more pronounced preference for intra-chromosomal interactions over inter-chromosomal interactions. Detection scores were calculated for each structure in each cell as follows:

### Chromosome Territories

The unnormalized Detection Score (DS) for chromatin territories is calculated using the following formula:

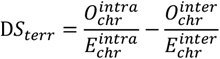

Here,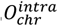 and 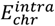 represent the observed and expected intra-chromosomal contact frequencies, respectively, while 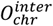 and 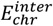 denote the observed and expected inter-chromosomal contact frequencies. The DS of chromatin territory is calculated in different chromtain interactions, such as chr1-chr2, and does not include the same chromosome interaction, such as chr1-chr1, with a contact map resolution of 100 Kb. Thus, taking detection score of the chr1 and chr2 as an example, 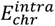 is calculated as 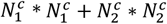,where 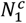 and 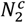 are the number of bins of chr1 and chr2. 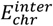 is calculated as 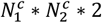. Finally, D*S*_*terr*_ of a single cell is the mean of the different chromatin interactions’ DS.

### A/B compartment

In contrast to scSPRITE, which requires a prior bulk A/B compartment structural data, we leverage the high-throughput capacity of sc3DG-seq data. By merging the top 300 cells with the most contacts from each dataset or the entire dataset, we generate a pseudobulk data. This data is used to identify A/B compartments, creating a prior compartment annotation for subsequent single-cell DS calculation. This approach avoids the need for prior bulk data, which may not always be available. The unnormalized DS of A/B compartments is calculated as follows:

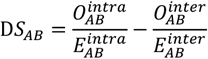

Here, 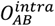 and 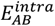 represent the observed and expected contact frequencies within compartments, respectively, while 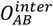 and 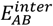 denote the observed and expected contact frequencies between compartments. The D*S*_*AB*_ of a cell is obtained by calculating the A/B compartment switches, which are transitions like ‘A-B-A’ or ‘B-A-B’, derived from the pseudobulk. Thus, taking a ‘*A*_1_-*B*_1_-*A*_2_’ with as an example, 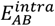 is calculated as 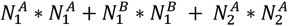 where 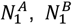 and 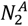 are the number of bins of *A*_1_ compartment, *B*_1_ compartment and *A*_2_ compartment. 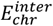 is calculated as 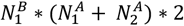. Finally, D*S*_*AB*_ of a single cell is the mean of all transition from pseudobulk. The calculation is performed on contact maps with a resolution of 100 Kb.

### Topologically Associating Domains (TADs)

Similar to the calculation of A/B compartments, the prior TAD structure is derived from merged cells. The formula for calculating the DS of TADs is as follows:

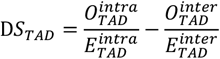

Here, 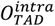 and 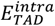 represent the observed and expected contact frequencies within TADs, respectively, while 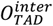 and 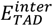 denote the observed and expected

contact frequencies between TADs. For the calculation of D*S*_*TAD*_, the 10 bins before and after the boundary are taken as two TADs. Thus, the calculation of D*S*_*TAD*_ is similar to the previous mentionded calculation method. The calculation is performed on contact maps with a resolution of 40 Kb.

Finally, we normalize the detection scores. For each structure in each cell, such as chr1-chr2 when calculating *DS*_*terr*_, ‘A-B-A’ or ‘B-A-B’ when calculating *DS*_*AB*_, and 10 bins before and after the detected boundary when calculating *DS*_*TAD*_, we sample the bins from the corresponding cells to construct the same structure and calcaulated the corresponding DS which repeat 100 times to get the mean of the results as the expected DS (Supplementary Fig.5a-c). Thus, the normalized detection scores(nDS) are calculated as the DS minus the expected DS.

### Insulation score and A/B compartment

To compute insulation score (IS) and annotate A/B compartments from single-cell or pseudobulk 3D genome data, we utilized the cooltools package (https://github.com/open2c/cooltools). IS, which measures the degree of chromatin insulation, was calculated using a contact map with a 40-Kb bin resolution. We used a window size that is ten times the bin size. Specifically, the ‘cooltools.insulation’ function was utilized with the parameter ‘ignore_diag=3’. A/B compartments were identified using a contact map with a 100-Kb resolution. The ‘cooltools.eigs_cis’ function was applied. We conducted a genome-wide computation without implementing specific filters to allow for a thorough analysis of genome compartmentalization. The parameter ‘n_eigs=3’ was used to capture the principal eigenvectors, which are indicative of the primary compartment patterns, with all other parameters remaining at their default settings.

### Accumulative single-cell quality analysis

For a comprehensive quality analysis of single-cell data, we implemented an iterative strategy to cumulatively aggregate individual cells based on either the top 300 cells with the highest contact counts or the entire dataset. This approach allows us to evaluate the consistency of 3D genome structure detection across cells within a sc3DG-seq dataset. To measure the similarity between the aggregated cells at each iteration and the final aggregate, we calculate the Intersection over Union (IoU) of the TAD boundaries. The IoU is calculated using the following formula:

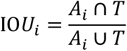

Here, *A*_*i*_ represents the set of TAD boundaries identified in the aggregated single-cell data at the *i*^*th*^ iteration, and T represents the set of TAD boundaries in the final aggregated cell. The IoU is a metric that quantifies the overlap between two sets of TAD boundaries. A higher IoU value signifies a greater degree of overlap, indicating higher consistency in the detected 3D structures.

## QUANTIFICATION AND STATISTICAL ANALYSIS

Statistical analyses were conducted using the Python 3.9.0 platform (https://www.python.org/)

## Data and code availability

STARK is freely available at https://github.com/CCCKW/stark. scNucleome is accessable at http://scnucleome.com/.

## Acknowledgments

This work was supported by the National Natural Science Foundation of China (32270683), the Beijing Natural Science Foundation (5242006), the Fundamental Research Funds for the Central Universities (BMU2021YJ064), the National Key R&D Program of China (2021YFC1712805), the National Natural Science Foundation of China (61572327, U20A20345 and 61972257), the CAMS Innovation Fund for Medical Sciences (2021-I2M5-003), the Natural Science Foundation of Shanghai (20JC1413800), and the National Key R&D Program of China (2020YFA0803803 and 2018YFA0900600). We gratefully acknowledge the High-performance Computing Platform of Peking University for conducting the data analyses.

## Author information

W.-J.J. and H.-J.W. conceived and designed the study. W.-J.J., and K.C. wrote the STARK software. W.-J.J., K.C., and Y.S. collected and analyzed the sc3DG-seq data with input from C.Z., R.G., N.W., F.L., A.L. and K.C. designed and constructed the scNucleome website. H.-J.W., M.X., X.Z., T.F., and Y.-J.W. supervised the study. W.-J.J., K.C., Y.S., and H.-J.W. wrote the manuscript with input from all authors. All authors reviewed and approved the final manuscript.

## Ethics declarations

The authors declare no competing interests.

