## supplemental File for "Harmonizing single cell 3D genome data with STARK and scNucleome"

**Supplementary Figures**

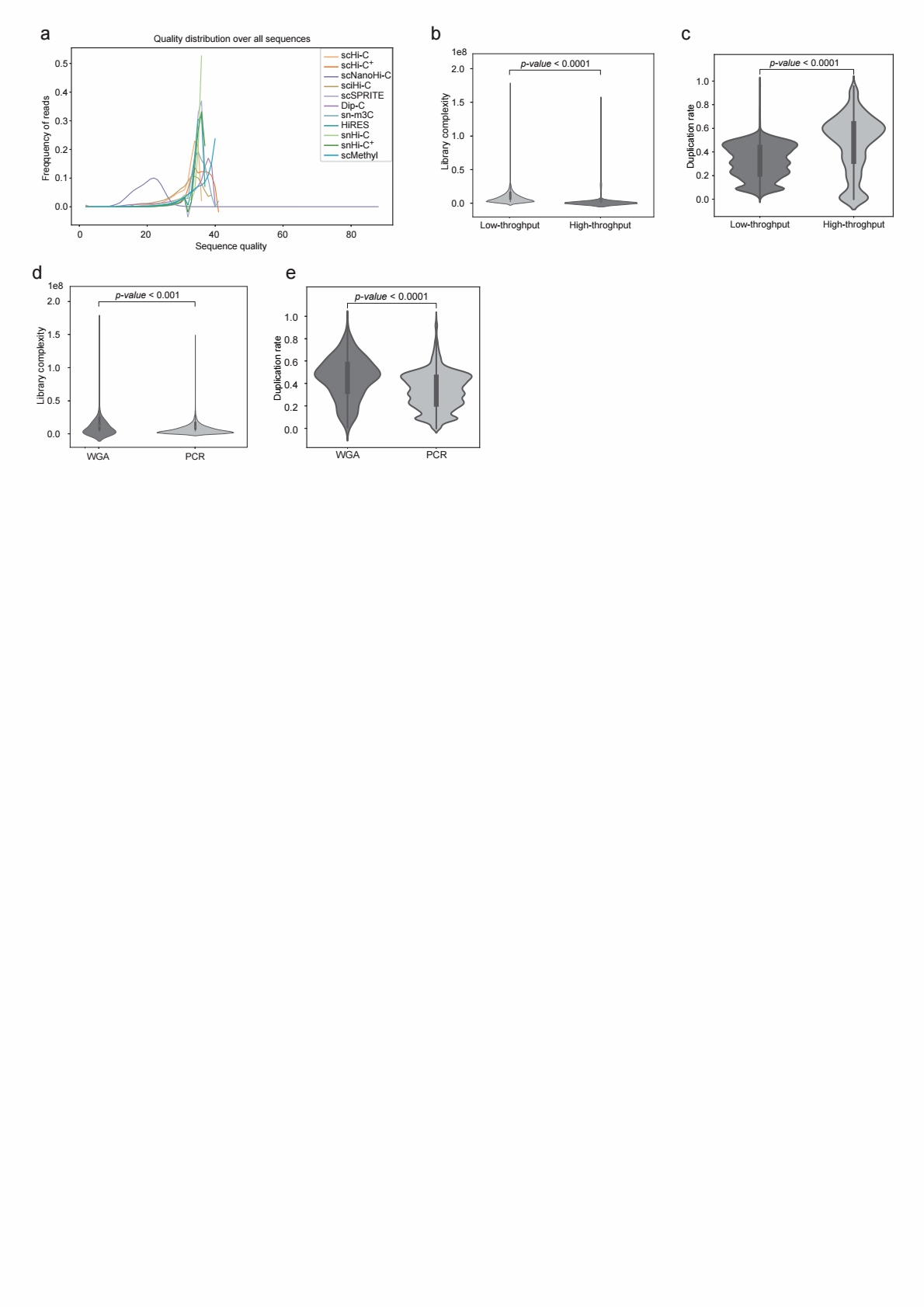

**Supplementary Fig.1** a) Line chart of the quality distribution of sequencing reads, library complexity and duplication rates across technologies; b) library complexity between high-throughput and low-throughput technologies; c) library complexity between technologies using PCR and those using WGA; d) Duplication rates between high-throughput and low-throughput technologies; e) Duplication rates between technologies using PCR and those using WGA. The Wilcoxon test p-value is shown in each figure.

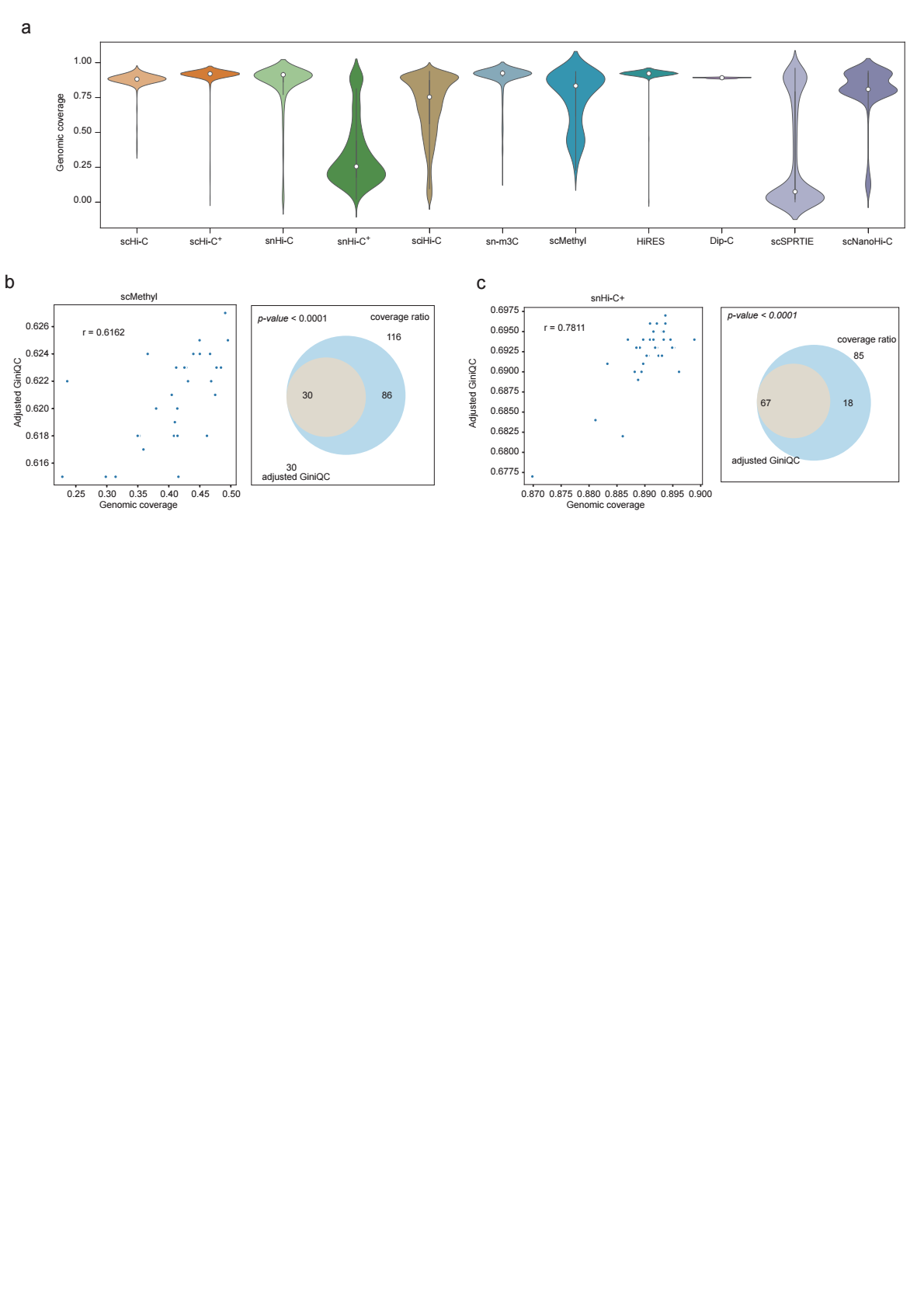

**Supplementary Fig.2** a) Analysis of genomic coverage and adjusted GiniQC across technologies; b) Left panel: Scatter plot of cells from scMethyl technology with adjusted GiniQC less than 0.65 and genomic coverage less than 0.5. Right panel: Venn diagram demonstrates the overlap of cells from scMethyl technology meeting both criteria. c) Left panel: Scatter plot of cells from snHi-C+ technology with adjusted GiniQC less than 0.65 and genomic coverage less than 0.7. Right panel: demonstrates the overlap of cells from snHi-C+ technology meeting both criteria.

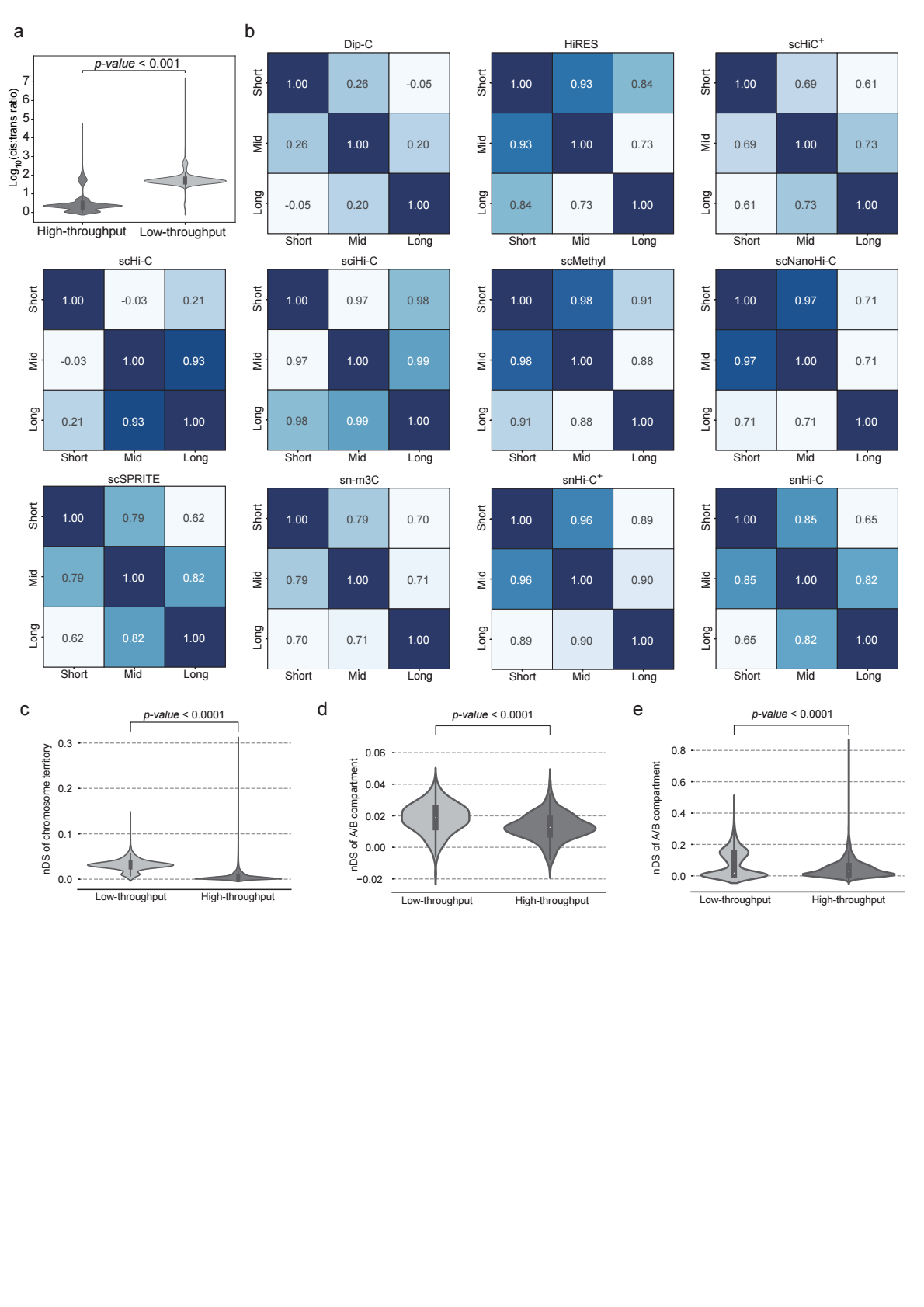

**Supplementary Fig.3** a) The cis:trans ratios between high-throughput and low-throughput technologies. b) Heatmap demonstrates the pair-wised correlation between contact counts across various genomic distance ranges for different technologies. c) Violin plot of nDS for chromatin territory between high-throughput and low-throughput technologies. d) Violin plot of nDS for A/B compartments between high-throughput and low-throughput technologies. e) Violin plot of nDS for TADs between high-throughput and low-throughput technologies. The Wilcoxon test p-value is shown in each figure.

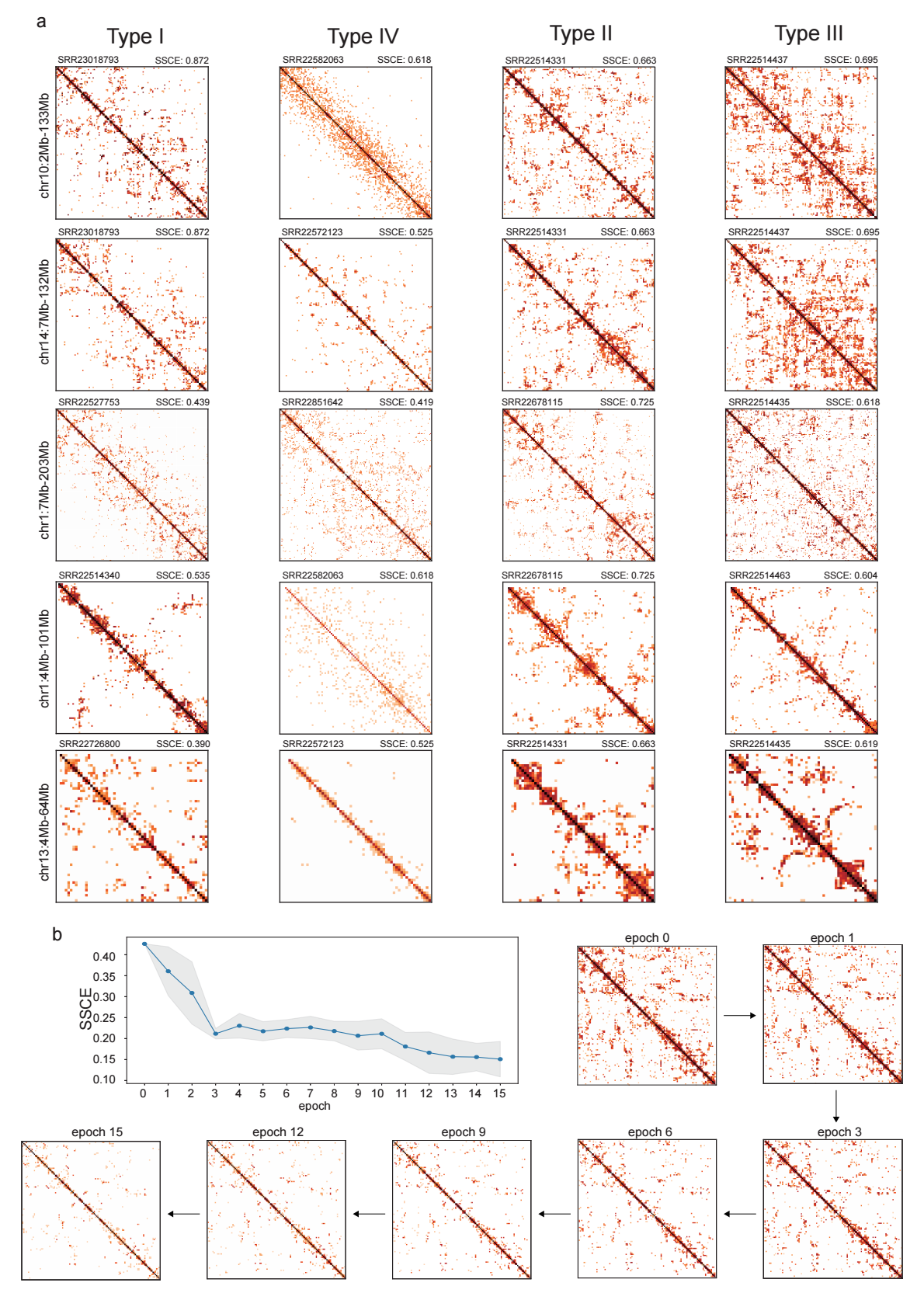

**Supplementary Fig.4** a) Visualization examples of contact maps for cells of four types in Fig. 4e. The cells displayed correspond to the red dots in the figure. Each row represents the contact map of the same genomic region across the four types. b) Line plot illustrates the change in Spatial Structure Capture Efficiency (SSCE) scores as the number of epochs increases. We simulate the gradual disappearance of genomic structures by continuously applying random masking to the contact matrix. In this simulation, we generate a masking matrix with the same dimensions as the contact matrix, with values uniformly sampled from the data range of the contact matrix. To address the possibility of obtaining negative values, which are not biologically meaningful in this context, we set all negative values to zero at each epoch. By repeating this process 15 times, we observe that SSCE values decrease as the number of epochs increases. This trend indicates that SSCE scores are sensitive to the decay of genomic structures, providing a measure of the integrity of chromatin interactions within the matrix.

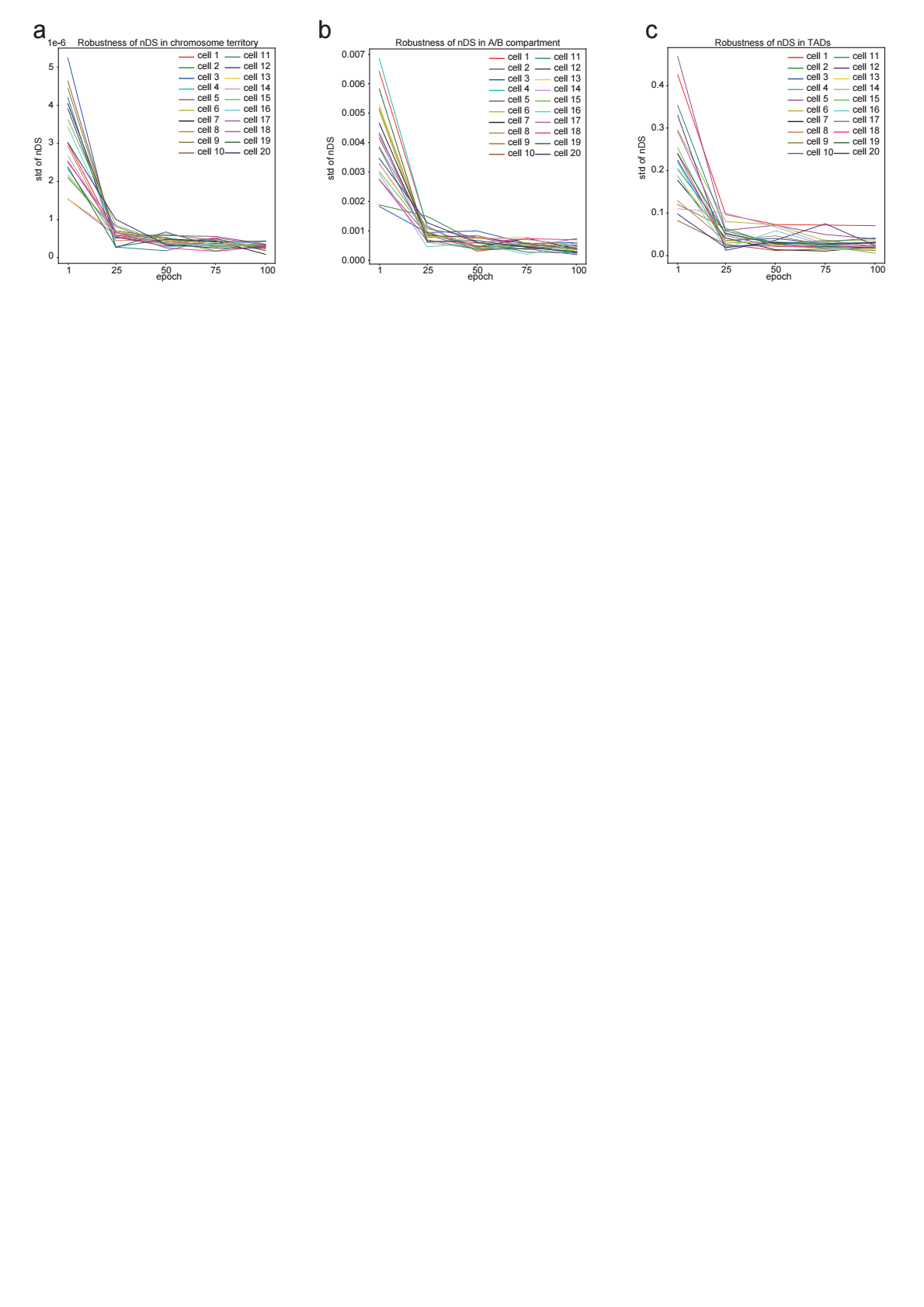

**Supplementary Fig.5**. Assessment of the robustness of the normalized detection score (nDS). Line plot indicates the influence of the number of epochs on the robustness of nDS calculations for different genomic structures: (a) chromatin territory, (b) A/B compartment and (c) TAD. The x-axis is the number of epochs, a hyper-parameter in the nDS calculation process. The y-axis displays the standard deviation of nDS values, which are derived from five independent calculations. The result reveals that the nDS values are stable when the number of epochs exceeds 25.
